# NMR assignments and secondary structure analysis of the eIF1-interacting fragment of human eIF3c

**DOI:** 10.1101/2025.09.13.675972

**Authors:** Simran Anand, Assen Marintchev

## Abstract

Eukaryotic translation initiation involves coordinated assembly of initiation factors on the small ribosomal subunit to form the pre -initiation complex (PIC) and scan mRNA for the start codon. The multi-subunit initiation factor eIF3 plays central roles in PIC assembly, stabilization, and scanning. Within eIF3c, residues 166–266 lie immediately N-terminal of the folded PINT/PCI domain and encompass the reported eIF1-binding site in human eIF3c. This segment is not visible in PIC Cryo-EM structures, except for a small helix contacting eIF1, and is predicted to be intrinsically disordered. Here we report the near-complete backbone and partial side-chain NMR assignments for the eIF3c 166–266 fragment in solution. The ^1^H–^15^N HSQC spectrum is consistent with an intrinsically disordered protein region. Several amide resonances were broadened at 25 °C but recovered at 5 °C. Secondary structure propensities derived from chemical shift index (CSI) analysis, together with amide ^1^H chemical shift temperature coefficients (CSTCs), confirm that the protein is disordered, while the CSI analysis also indicates the presence of short segments with modest α-helical or β-strand propensity. Three conserved FLKK motifs fall at junctions of these transient structural elements, with Motif 3 located in the subsegment showing slightly greater propensity for transient structure. These residue-specific NMR assignments provide a foundation for future studies of interaction surfaces, binding-induced folding, and conformational dynamics of this conserved eIF3c region in the context of translation initiation.

## Biological Context

Eukaryotic translation initiation is a multi-step process in which a functional ribosome is assembled on an mRNA to direct protein synthesis. A pivotal stage is the formation of the pre-initiation complex (PIC), where the small ribosomal subunit associates with eukaryotic translation initiation factors (eIFs) including eIF1, eIF3, eIF5, and eIF2 and the initiator Met-tRNAi, to scan the mRNA for a start codon in an appropriate sequence context. Within this network, eIF3 plays a central role in PIC assembly and stabilization, and scanning (reviewed in (Brito Querido et al., 2024, Dever et al., 2023, Hinnebusch, 2014, Marintchev, 2025)).

eIF3c residues 166–266 are positioned immediately upstream of the folded PINT/PCI domain, which anchors eIF3c within the eIF3 scaffold and interacts with the 40S subunit and the mRNA exit channel (reviewed in (Valasek et al., 2017)). In human eIF3c, this region contains the binding site for eIF1, reported to be located between residues 166–287 (Brito Querido et al., 2020), 171-240 (Bochler et al., 2020), or 165–213 (Kratzat et al., 2021). In contrast, in *Saccharomyces cerevisiae (S. cerevisiae)*, eIF1 binds to the disordered N-terminal region preceding the small folded domain of eIF3c (Karaskova et al., 2012, Obayashi et al., 2017), with reported additional interactions involving the folded domain in some studies (Obayashi et al., 2017).

In contrast to the PCI domain, the 166–266 region is not visible in Cryo-EM structures, except for a short helix bound to eIF1, and is predicted to be intrinsically disordered (Brito Querido et al., 2020, des Georges et al., 2015, Kratzat et al., 2021). The low local resolution has not allowed identifying which amino acids in human eIF3c comprise the eIF1-binding helix. Such intrinsic disorder is often associated with conformational plasticity, allowing a single sequence to engage multiple molecular partners and adapt to structural rearrangements during initiation. Highly charged disordered segments in other eIF3 subunits are likewise implicated in dynamic, multi-partner interactions within the PIC (reviewed in (Valasek et al., 2017)). The 166–266 region of eIF3c is highly conserved across *Metazoa* species, engages in molecular interactions with other PIC components, and displays structural flexibility (Petrychenko et al., 2024). These features suggest it is a key functional module within eIF3, and detailed characterization of this segment would address an important missing piece in our molecular understanding of the complex.

Despite biochemical evidence for the functional significance of the 166–266 fragment, its structural properties and precise interaction surfaces have not been determined. This lack of atomic-level detail limits our understanding of how eIF3c regulates the PIC architecture and initiation factor dynamics. NMR spectroscopy is particularly well suited to define the structural propensities and dynamics of such intrinsically disordered yet interaction-prone regions (reviewed in (Dyson & Wright, 2004, Marintchev et al., 2007)).

## Methods and Experiments

### Protein expression and purification

The codon-optimized DNA sequence for human eIF3c-166-266 was cloned into a pET21a (Novagen) derivative plasmid with an N-terminal GB1 tag (the IgG-binding domain 1 of *Staphilococcus aureus* protein G), followed by a His_6_ tag and a TEV protease cleavage site, and a C-terminal cysteine.

Recombinant GB1-tagged eIF3c-166-266 protein was expressed in *Escherichia coli BL21(DE3)* cells. The expression plasmid was transformed into chemically competent cells and plated on LB agar containing carbenicillin. A single colony was used to inoculate 15 mL LB starter cultures, supplemented with carbenicillin, and grown overnight at 37 °C. The following day, 1 L minimal medium containing ^15^NH_4_Cl and/or ^13^C-glucose was inoculated to an initial OD_600_ of 0.05–0.15. Cultures were grown at 37 °C with shaking until OD_600_ reached 0.6–0.8, then induced with 1 mM IPTG. Expression proceeded for 2–3 h at 37 °C or overnight at 20 °C.

Cell pellets from 1 L cultures were resuspended in 40 mL lysis buffer containing 10 mM sodium phosphate (pH 7.0), 300 mM KCl, 7 mM β-mercaptoethanol, 0.1 mM AEBSF, protease inhibitor cocktail, 1 mg/mL lysozyme, and 0.01% NaN_3_. Cells were lysed by sonication and the lysate clarified by centrifugation. Soluble protein was bound to pre-equilibrated Talon CellThru resin by batch incubation at 4 °C, washed extensively with running buffer (300 mM KCl, 10 mM sodium phosphate, pH 7.0, 7 mM β-mercaptoethanol, 0.1 mM AEBSF, and 0.01% NaN_3_), and running buffer with 4 mM imidazole, and eluted with running buffer containing 200 mM imidazole. Elution fractions were supplemented with 2 mM DTT and 1 mM EDTA.

Ion-exchange chromatography purification was performed on a HiTrap Q FF 5 mL column, using a low-salt buffer containing 100 mM NaCl, 10 mM sodium phosphate, pH 7.0, 1 mM DTT, 1 mM EDTA, 0.1 mM AEBSF, and 0.01% NaN_3_, and a high-salt buffer containing 1 M NaCl, 10 mM sodium phosphate, pH 7.0, 1 mM DTT, 1 mM EDTA, 0.1 mM AEBSF, and 0.01% NaN_3_ for gradient elution. The column was equilibrated in the low-salt buffer. Protein separation was achieved using a 25 mL gradient from 0% to 50% high-salt buffer.

The GB1 tag was cleaved using TEV protease in buffer containing 150 mM NaCl, 10 mM sodium phosphate, pH 7.0, 1 mM DTT, and 1 mM EDTA at a ratio of 1:50 for two hours at room temperature, leaving an N-terminal glycine from the tag. The tag and TEV were then removed by ion-exchange chromatography, as described above.

### NMR Spectroscopy

The protein samples were exchanged in NMR buffer, containing 10 mM sodium phosphate, pH 7.0, 150 mM KCl, 2 mM DTT, 1 mM EDTA, and 5% D_2_O. Triple-resonance NMR spectra for backbone assignments, including HNCO, HN(CA)CO, CBCAHN, CBCA(CO)NH, and (H)C(CO)NH, and ^15^N-TOCSY, were performed on a Bruker AMX 500 MHz, equipped with a cryoprobe at both 25 °C and 5 °C. Two temperatures were used because a number of ^1^H–^15^N HSQC peaks were missing or exhibited line broadening at 25 °C, and to allow analysis of temperature-dependent chemical shift changes. ^1^H–^15^N HSQC spectra were acquired at both temperatures, and proton chemical shift changes between the two conditions were examined.

All NMR data were processed using NMRPipe (Delaglio et al., 1995), and backbone assignments were completed using CARA (Keller, 2004).

## Extent of assignments and data deposition

The ^1^H–^15^N HSQC spectrum for the human eIF3c 166–266 fragment is shown in Figure 1. The spectrum displays poorly resolved peaks consistent with an intrinsically disordered protein segment, with missing or weak resonances corresponding to residues broadened by conformational exchange or exchange with the water. Backbone resonance assignments are near complete, with ^1^H–^15^N amide correlations assigned for 95/99 non-proline residues, corresponding to 96.0 %. Four non-proline residues could not be assigned (Ser 166, Glu 229, Glu 265, and Asp 266). Several NH peaks were very weak or missing at 25 °C but became visible at 5 °C, aiding unambiguous identification of these residues — for example, A167, S201, S205, S208, G209, S235, S237, S238, S254, along with other residues previously broadened at 25 °C, could be clearly observed at the lower temperature.

**Figure 1.**
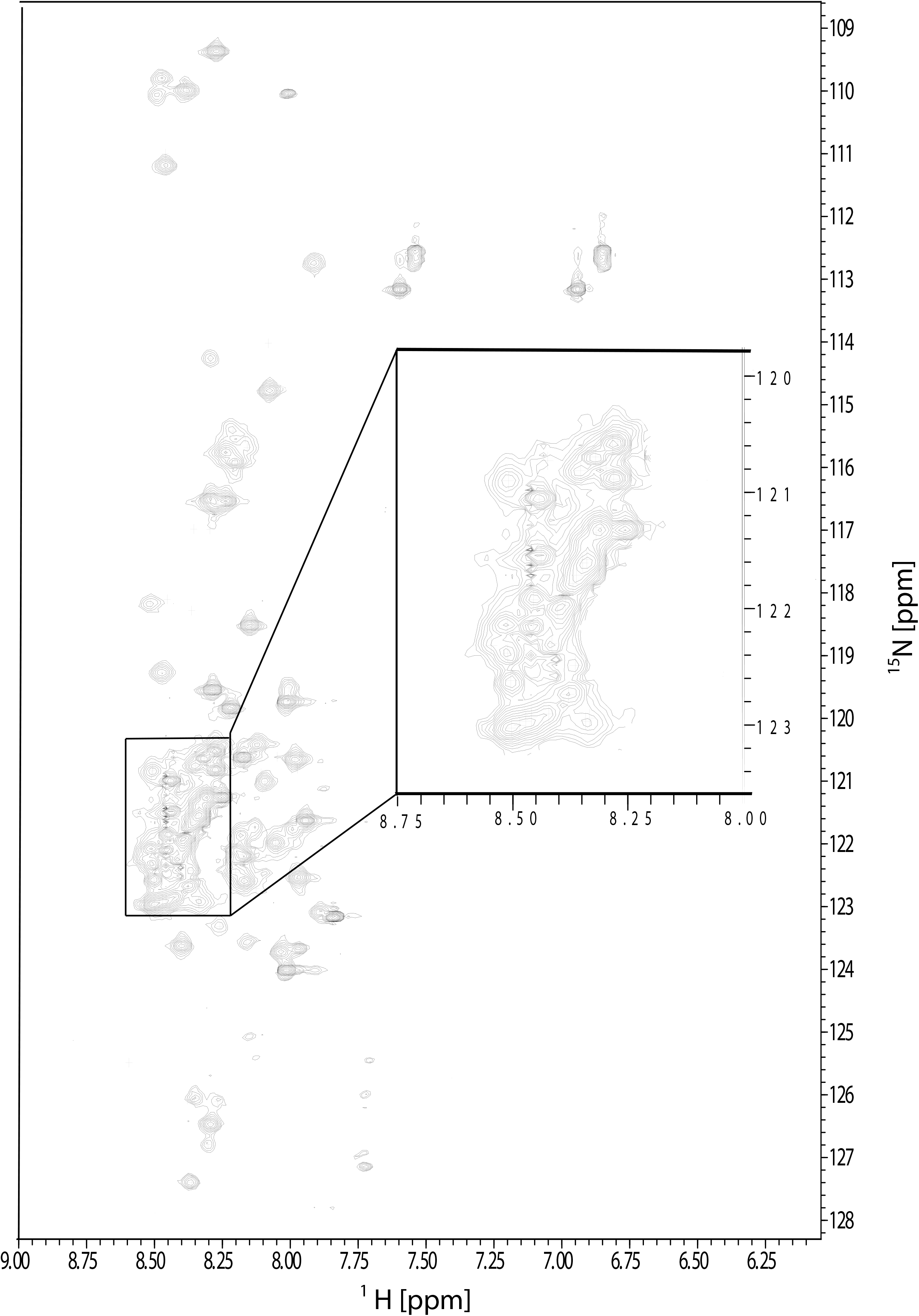
Annotated 2D ^1^H, ^15^N-HSQC spectra for the human eIF3c 166-266 fragment. Spectra were recorded at 298 K at a field strength of 500 MHz, in a buffer containing 10 mM sodium phosphate, pH 7.0, 150 mM KCl, 2 mM DTT, 1 mM EDTA, and 5% D_2_O. Peaks corresponding to degradation products are denoted with black squares. The limited dispersion of backbone amide resonances is characteristic of an intrinsically disordered protein.

Cα resonances are assigned for 95/101 residues (94.1 %). Cβ resonances are assigned for 90/96 expected residues (93.8 %). Beyond the Cβ, assignments for side-chain carbons observed in the (H)C(CO)NH spectra (excluding aromatic rings, the methyl carbon of methionines, and the Cζ carbon of arginines) were completed for 37/47 expected residues. Carbonyl (CO) carbons are assigned for 69 / 101 expected residues (68.3 %).

The ^1^H, ^13^C, and ^15^N NMR chemical shift assignments for the human eIF3c (166-266) fragment have been deposited in the Biological Magnetic Resonance Data Bank (https://bmrb.io/) under accession code 53212.

### Analysis of secondary structure

Secondary structure propensities were evaluated using two complementary NMR-based approaches (Figure 2). The first was chemical shift index (CSI) analysis, using the CSI 3.0 server (Hafsa et al., 2015), This approach identifies α-helical and β-strand regions by comparing experimentally observed backbone NMR chemical shifts to a set of residue-specific ‘random coil’ chemical shifts. These random coil values serve as a baseline, representing the expected shifts for an unstructured polypeptide. For this analysis, the backbone chemical shifts of ^1^Hα, ^13^Cα, ^13^Cβ, and ^13^CO nuclei were used. Based on this comparison, the server converts significant downfield or upfield secondary shifts into simple integer scores: +1 for α-helical propensity, –1 for β-strand propensity, or 0 for random coil. A significant secondary structure element is identified by a consensus of these scores over a stretch of amino acids. A consensus of four or more consecutive residues with CSI values of +1 is a robust indicator of an α-helix, while a consensus of three or more consecutive residues with values of –1 indicates a β-strand. This method provides a well-established and rapid measure of local secondary structure bias (Hafsa et al., 2015).

**Figure 2.**
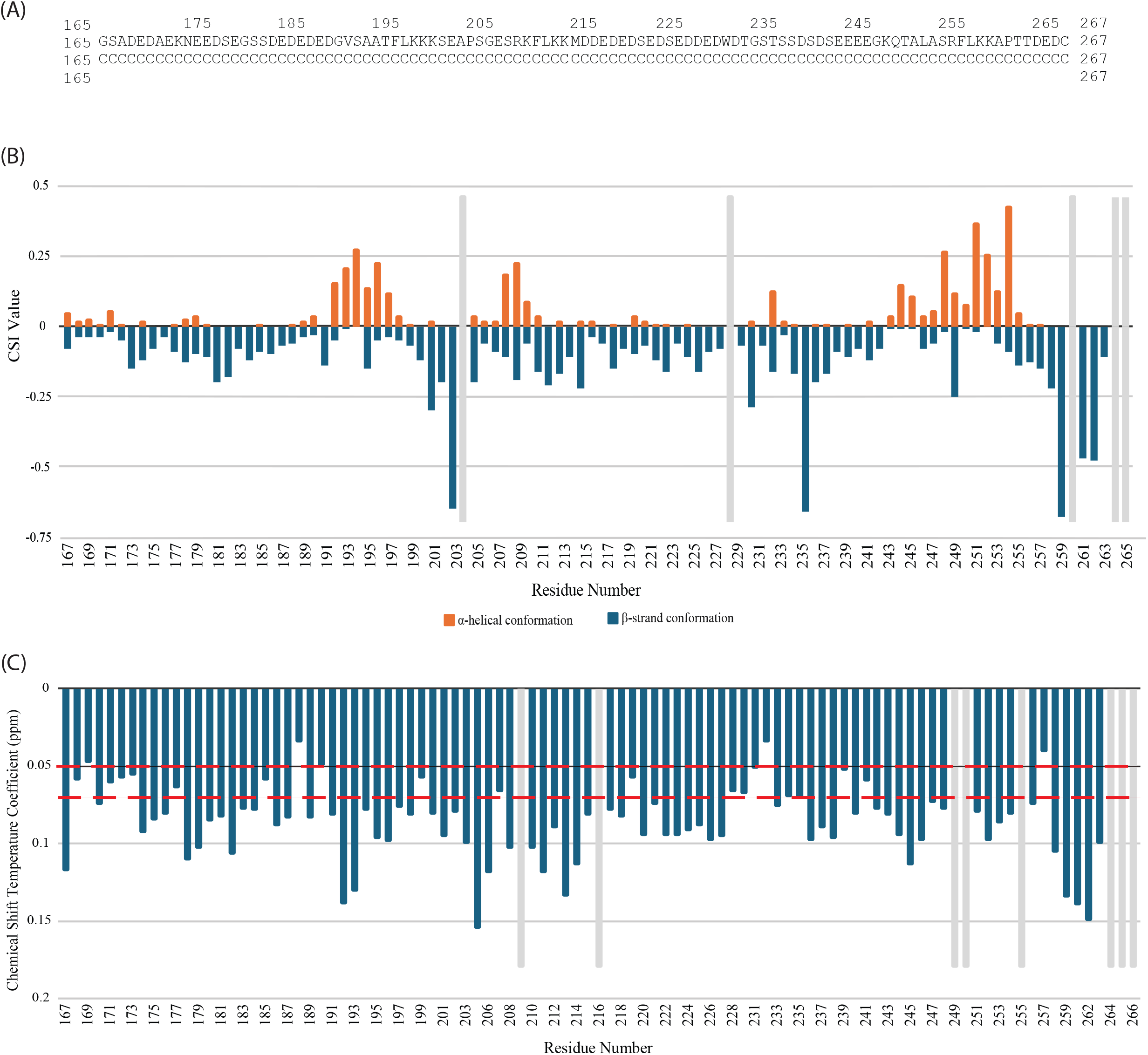
NMR-based secondary structure analysis of eIF3c residues 166–266. The secondary structure propensity of the eIF3c segment was evaluated using Chemical Shift Index (CSI) and amide ^1^H chemical shift temperature coefficient (CSTC) analyses. (A)Primary sequence and secondary structure prediction of the eIF3c 166-266 fragment from CSI analysis. The 103-amino acid sequence of the studied segment (human eIF3c residues 166-266 with an N-terminal glycine and C-terminal cysteine) is shown. The secondary structure, determined from Chemical Shift Index (CSI) data, is assigned as random coil (‘C’) for the entire region. The sequence contains three conserved FLKK motifs, located at residues 196–199, 211–214, and 256–259. (B) Chemical Shift Index (CSI) analysis. CSI values were calculated using the CSI 3.0 server. Values α-helix (orange bars) and β-strand (blue bars) are plotted for each residue. Unassigned residues are indicated by light grey bars. (C) Amide ^1^H chemical shift temperature coefficients (CSTCs). Statistical reference lines are shown at –3.6 ppb/K (dotted line) and –2.5 ppb/K (dashed line), corresponding to one and two standard deviations, respectively, above the established mean for intrinsically disordered regions (Okazaki et al., 2018). Critically, no strings of consecutive residues exhibit a CSTC value less negative than the two-standard-deviation threshold, confirming the absence of stable, hydrogen-bonded secondary structure in this segment. Unanalyzable residues are indicated by light grey bars.

The second was analysis of amide ^1^H chemical shift temperature dependence, in which chemical shift temperature coefficients (CSTCs) are determined from the slope of the resonance frequency change versus temperature. Smaller-magnitude (less negative) CSTC values correlate with amides that are buried and/or hydrogen-bonded, thus likely folded. While this analysis may not distinguish between an amino acid in an intrinsically disordered region (IDR) and one in a folded region that is solvent-exposed and not hydrogen-bonded, it can identify residues that are likely to be hydrogen-bonded at least transiently (Cierpicki et al., 2002, Okazaki et al., 2018). Therefore, we focused on resonances that experience much smaller changes than expected for a fully disordered region. Prior hydrogen-bond correlation work in ubiquitin suggested that residues with less negative slopes (≥ –4.6 ppb/K) are typically hydrogen-bonded, and those with more negative slopes (< –4.6 ppb/K) are solvent-exposed (Cierpicki et al., 2002). Okazaki and coauthors found that disordered residues averaged –5.62 ± 1.56 ppb/K, whereas ordered residues averaged –3.19 ± 2.58 ppb/K and further optimized the discrimination of IDR/non-IDR residues at a –3.6 ppb/K cutoff (Okazaki et al., 2018). Using the data from Okazaki and co-authors (Okazaki et al., 2018), we applied a cutoff at the disordered region average plus two standard deviations (mean + 2 SD = –2.5 ppb/K); residues with CSTCs less negative than this threshold have an increased probability of not being fully disordered.

The combined CSI and CSTC analyses revealed that this region of eIF3c is intrinsically disordered overall (Figure 2). CSI indicated modest propensity for helical conformation in residues 192–197, 208–211, and 244–256, and for extended (β-strand-like) conformation in residues 200–206, 211–214, and 258–264 (Figure 2B), none of which exceeded the CSI 3.0 server threshold to be classified as secondary structure. These tendencies are likely transient, meaning the segment probably spends only a fraction of the time in helical or extended conformations — a dynamic behavior commonly associated with intrinsically disordered regions that mediate molecular recognition. The CSTC analysis confirms that this region is intrinsically disordered, as most measured CSTC values fall within the expected range for a disordered polypeptide, with only a few isolated residues exhibiting values outside the -2.5 ppb/K cutoff (Figure 2C).

The three conserved FLKK motifs fall within or immediately at the boundary between CSI-defined modest helix and strand propensities: Motif 1 (196–199) lies at the junction between segments 192–197 and 200–206, Motif 2 (211–214) coincides with the 211-214 segment with slight strand propensity, and Motif 3 (256–259) lies at the junction between segments 244–256 and 258–264. All three occur in zones of above-average secondary structure propensity for this disordered segment, and Motif 3 lies within the sub-segment with the highest propensity (Figure 2B).

## Conclusion

Here, we report the backbone NMR chemical shift assignments for the human eIF3c-166-266 fragment, in order to facilitate future studies exploring at an atomic level its interactions and dynamics.

The combined chemical shift index (CSI) and chemical shift temperature coefficient (CSTC) analyses demonstrate that the 166–266 region of human eIF3c is predominantly intrinsically disordered under physiological conditions, but harbors distinct subsegments with modest propensities for α-helical or β-strand secondary structure. The CSI profiles identify three short stretches with helical bias, each followed by a stretch with extended/β-strand-like bias (Figure 2B). Together, these findings are consistent with a dynamic conformational ensemble in which transient secondary structure formation occurs in localized sequence regions embedded within an otherwise flexible chain. Notably, a portion of the human eIF3c 166-266 segment forms a short helix upon binding to eIF1 (Brito Querido et al., 2020, Kratzat et al., 2021, Petrychenko et al., 2024). The observation that even the highest secondary structure propensities are rather small underscores the fundamentally disordered nature of this region.

This is in line with bioinformatic predictions and the absence of consistent cryo-EM density for this segment in PIC structures (Brito Querido et al., 2020, Kratzat et al., 2021, Petrychenko et al., 2024).

Remarkably, all three conserved FLKK motifs are positioned at or near junctions between CSI-defined helical and extended-bias regions. This arrangement could be functionally significant: junctions between local structural biases may act as adaptable hinges or recognition sites in disordered binding regions, allowing context-dependent shifts in conformation and presentation of key side chains. Motif 3 (256–259) resides within the subsegment with the highest CSI indicators of transient structure, suggesting that it may sample a folded state more frequently even in the absence of binding partners. This observation suggests that Motif 3 could be involved in the interaction with eIF1.

Overall, the NMR resonance assignments and analysis presented here establishes a detailed baseline for future functional and structural studies of eIF3c 166–266. These assignments and propensity maps enable future high-resolution mapping of binding-induced folding events, and integration with cryo-EM interaction data to build a more complete picture of eIF3 architecture and function during translation initiation.

The combined analysis reveals the region is predominantly disordered. However, CSI identifies segments with modest propensities for α-helical conformation (residues 192–197, 208–211, and 244–256) and β-strand conformation (residues 200–206, 211–214, and 258–264).

## Statements and Declarations

## Acknowledgements

This work was supported by the National Institute of Health [GM134113 to A.M.]. We thank the NMR Core Facility at Boston University Chobanian & Avedisian School of Medicine for access to instrumentation and technical support.

## Author Contributions

S.A. expressed and purified the protein construct, prepared isotopically labeled samples, performed NMR data analysis, assigned resonances, and conducted secondary structure analysis.. A.M. collected NMR spectra, guided experimental design and supervised all stages of the project. S.A. and A.M. jointly interpreted the data and wrote the manuscript. All authors reviewed and approved the final version.

## Data Availability

The chemical shift assignments have been deposited in the Biological Magnetic Resonance Data Bank (BMRB) under accession number **53212**.

## Ethical Approval

The authors declare that no human or animal subjects were involved, and all experiments were conducted in compliance with institutional and ethical guidelines.

